# Simvastatin and Primaquine Identified as Potential Endometriosis Therapeutics via a Novel Epithelial-Stromal Assembloid Drug Screening Assay

**DOI:** 10.64898/2026.09.27.754752

**Authors:** Ferheen Abbasi, Xinyu Tang, Juan C. Irwin, Binya Liu, Tomiko T. Oskotsky, Marina Sirota, Frederick J. Meyers, Linda C. Giudice

**Affiliations:** Department of Obstetrics, Gynecology & Reproductive Sciences, UCSF, San Francisco, CA, USA; School of Medicine, UC Davis, Sacramento, CA, USA; Mass General Brigham Hospital, Harvard University, Boston, MA, USA; Bakar Computational Health Sciences Institute, UCSF, San Francisco, CA, USA; Department of Pediatrics, UCSF, San Francisco, CA, USA; Division of Clinical Informatics and Digital Transformation, Department of Medicine, UCSF, San Francisco, CA, USA; Department of Internal Medicine, Division of Hematology and Oncology, UC Davis Health, Sacramento, CA, USA

**Keywords:** Endometriosis, endometrium, assembloids, therapeutics, transcriptomics, drug repositioning

## Abstract

**Background:** Endometriosis is an estrogen-driven, inflammatory disorder affecting ∼10% of menstruators, causing severe pain. Current treatments that reduce estrogen or inflammation have inconsistent efficacy and poor tolerability. Therapies identified via drug-repositioning methods, which target patient-specific pathways, offer a promising alternative. Simvastatin and primaquine were identified as potential treatments, strongly reversing endometriosis-associated gene expression pathways, and behavioral testing in an animal model showed that these drugs diminish endometriosis-associated pain. This paper examines their effects in human assembloids to provide proof-of-concept validation and insights into their mechanisms of action relevant to endometriosis.

**Methods:** We established human endometrial assembloids using immortalized epithelial (12Z) and stromal fibroblast (iEc) cell lines, as well as primary tissue from patients. Assembloids were generated in 96-well agarose molds with both cell types. We tested ibuprofen, fenoprofen, primaquine, and simvastatin. Following a 24-hour exposure, assembloids and monolayer cultures were harvested and stranded mRNA-seq libraries were sequenced on an Illumina NovaSeqX Plus System.

**Results:** Gene set enrichment analysis showed key pathway reversals: Primaquine reversed the chemical carcinogenesis ROS pathway in assembloids. Simvastatin reversed cytokine-cytokine receptor interaction in both epithelial and stromal cell lines, and the cytoskeleton in muscle cells pathway in stromal cell lines. Fenoprofen reversed the calcium signaling pathway in stromal cell lines.

**Conclusion:** Simvastatin induced the most significant gene expression changes, notably reversing cytokine–cytokine receptor interactions in both cell lines, mirroring findings from our rat model. Despite the small sample size limitation, these experiments highlight the promise of assembloid models to test therapeutic candidates for endometriosis.

## Introduction

Endometriosis is a common, inflammatory, estrogen-driven disorder wherein endometrial tissue, attaches and invades pelvic structures, eliciting inflammation, fibrosis, and pain, and infertility and poor pregnancy outcomes. Diagnosis relies on surgical and histologic confirmation of disease and pelvic imaging.[1] Medical management includes nonsteroidal anti-inflammatory drugs (NSAIDs), analgesics, neuromodulatory drugs and hormone therapies[2,3] Notably, 25- 34% of patients treated medically have no decrease in their endometriosis-associated pain symptoms and in 60%, 5-16% of patients abandon hormonal therapies due to intolerable side effects, and symptoms recur within 2 years; surgical excision or ablation demonstrate similar results.[2] Thus, there is a huge unmet need for personalized and targeted therapies.[4–6]

Computational drug repurposing aids in the quest for treatment by uncovering novel therapeutics for existing, FDA-approved drugs beyond their original clinical indication.[7–10] Our group previously developed a transcriptomics-based strategy based on the hypothesis that effective therapeutics can counteract disease-associated gene expression patterns. Under this framework, genes down-regulated in the disease state are up-regulated by the drug, and vice versa[11] This approach has been successfully identified candidate therapies across a range of diseases, including inflammatory bowel disease,[12] preterm birth,[13] Alzheimer’s disease[14] and, recently, endometriosis.[15,16] In the latter study, publicly available microarray data from eutopic endometrial tissue from subjects with and without endometriosis identified fenoprofen, primaquine, and simvastatin as potential therapeutic candidates for endometriosis.[15,16] This drug repurposing study identified 299 compounds predicted to reverse endometriosis-associated gene expression profiles leveraging endometriosis transcriptional signatures both without stratification and stratified by ASRM disease stage (I–II vs. III–IV) and menstrual cycle phase (proliferative, early secretory, and mid-secretory). Among the top candidates, fenoprofen and simvastatin significantly reduced vaginal hyperalgesia in a rat model of endometriosis, whereas primaquine exhibited comparatively modest analgesic effects. [15,16]

The current study aims to determine the effects of simvastatin and primaquine on the transcriptomes and signaling and biological pathways of endometrial, stromal, and epithelial cells. Specifically, we established human endometrial assembloids comprised of two immortalized epithelial (12Z) and stromal (iEc) endometriosis cell lines, and primary cell lines from eutopic endometrial tissue from patients with endometriosis. Based on their strong reversal of the endometrial bulk RNA sequencing signatures *in silico* and in our *in vivo* rat model, and our recent analysis of electronic medical records showing that simvastatin was associated with reduced risk of endometriosis diagnosis, we treated the assembloids with simvastatin, primaquine, and fenoprofen, with ibuprofen as a control.[15,16] In the current study, bulk RNA sequencing revealed that simvastatin and primaquine reversed numerous pathways involved in inflammation in human endometrial assembloids, and fenoprofen reversed calcium signaling in stromal cells. These data support the value of combining computational drug repurposing with endometrial assembloid validation to identify novel, non-hormonal therapies for processes associated with endometriosis inflammation and pain.

## Methods

### Collection of Endometrial Samples

Human eutopic (within the uterus) endometrial tissue samples were collected through the UCSF/NIH Human Endometrial Tissue/DNA Bank from premenopausal women undergoing gynecological procedures at University of California, San Francisco according to an International Review Board-approved protocol (IRB#10-02786). Patients using hormonal birth control were excluded. Endometriosis patients were surgically confirmed and had no other gynecologic abnormalities.

## Cell Culture

Three cell lines were used in the experiments: immortalized human endometriotic epithelial cells (12Z), human endometrial fibroblast ETB 128++ cells, and immortalized human endometriotic stromal cells (iEc-ESC). The iEc-ESC cells were a generous gift of Dr. Azgi Fazleabas.[17] After thawing at 37℃, approximately 1 mL of cells (10^6 viable cells count) was transferred to a 50 mL tube. Cells were washed with 9 mL of DMEM + Glutamax (Thermo Fisher Scientific, 11594446) supplemented with 10% FBS and penicillin-streptomycin, added dropwise with gentle agitation, and then centrifuged at 300 x g for 5 minutes. The supernatant was removed and 1 mL DMEM was added. After counting, the cells were plated in 10 cm Petri dishes containing 9 mL DMEM. The medium was changed every 2-3 days. 12Z and ETB cells were harvested after 4-5 days, whereas iEc-ESC cells were harvested after 7-8 days.

## Cell Line Assembloid Creation

Cell line assembloids were created via an adapted protocol.[18] First, micro-molds (MicroTissues 3D Petri Dish) were created using the published protocol. Briefly, 330 uL of autoclaved agarose in 0.9% NaCl was added to the 96-well micro-mold, refrigerated for 5 minutes, and flexed out gently using forceps. The agarose mold was transferred to a 24 well plate in 1 mL of MammoCultTM Human Medium (Stemcell Technologies, 05620) supplemented with hydrocortisone and heparin according to the manufacturer’s instructions. The molds were incubated for 15 minutes to equilibrate with culture medium and then the media was changed twice. The molds were kept in the incubator for a maximum of 1 week. Next, endometrial cell line monolayer cultures were harvested by trypsinization, centrifuged 300xg for 5 min, and resuspended with 1 mL of Mammocult. The viable cell concentration was adjusted to 1.2M cells/mL, and ∼60,000 cells/50 uL were seeded into 96-well agarose molds with the epithelial to stromal ratios of: 1:1, 1:2,1:5, 1:10, 1:50, 2:1, 5:1, 10:1. Assembloids were allowed to settle without media change for four days, and then fresh media was supplemented every other day. On the 8^th^ day, assembloids were evaluated for number and size distribution on a Zeiss Axio Observer Z1 inverted microscope equipped with bright field and phase-contrast optics, using Zen imaging software. Subsequently, assembloids were processed for histological evaluation. Briefly, assembloids were fixed in 4% paraformaldehyde in phosphate buffered saline, dehydrated, and paraffin embedded. Four µm sections were deparaffinized and stained with hematoxylin and eosin (H&E).

## Primary Endometrial Tissue Assembloid Creation

Epithelial and stromal cells were isolated from endometrial tissue by collagenase digestion and size fractionation as previously described.[19] Cryopreserved tissue was thawed, minced, and digested with collagenase and Dnase for 1-2 hours at 37C with gentle agitation. The tissue digest passed first through a 100 μm cell sieve to remove undigested tissue, and the flowthrough then through a 20 μm cell sieve to separate the epithelial glands from single stromal cells.

Contaminant erythrocytes in the stromal fraction were removed by incubation with 00-4333-57 (ThermoFisher 00-4333-57). After centrifuging and resuspending with assembloid media, stromal cells were counted and adjusted to 1.2 M cells/mL. The epithelial cell fraction was adjusted to roughly the same density as the stromal using microscopy as described in the original protocol,^18^ and seeded at following ratios of epithelium to stroma (E:S): 3:1, 1:5, 1:10, 1:20, and cultured as above for the cell lines. After the 8th day in culture, assembloids were either fixed for H&E staining as described above for the cell line organoids, or drug tested.

## Drug treatment of assembloids

Four compounds of interest were chosen for drug treatment trial: ibuprofen (IBU), fenoprofen (FENO), simvastatin (SIM), and primaquine (PRIM). Concentrations for each were selected based on the Connectivity MAP data (IBU: 19.4 uM; FENO: 7.2 uM; SIM: 9.6 uM; PRIM: 8.8 uM) to validate the assembloid model. Drugs were dissolved in DMSO. Cell line assembloids were created using the 1:50 ratio of epithelial 12Z to stromal iEc cells. We allocated 4 wells/ condition (∼240,000 cells total), with the 4 drugs and vehicle control (DMSO). After 1 week of growth and a 24-hr drug test, assembloids were harvested, pooled per condition, RNA extracted (Qiagen Mini RNeasy Kit), and submitted for bulk RNA sequencing. The same drug testing protocol was repeated for primary tissue assembloids from 2 patients with endometriosis.

## RNA isolation

RNA was isolated from cultured cells using Trizol (Thermofisher) and further purified using the RNeasy kit (Qiagen) with the optional on-column DNase step according to the manufacturers’ protocols. Total RNA was eluted from the columns in nuclease-free water and stored at -80°C.

RNA concentration and purity were assessed with a NanoDrop 2000 Spectrophotometer (Thermo Scientific) and quality assessments (e.g., RNA integrity) were made using an Agilent 2100 Bioanalyzer (Agilent Technologies).

### Preparation of directional RNA-sequencing (RNA-seq) libraries and data processing

Indexed, stranded mRNA-seq libraries were prepared from total RNA (300 ng) using the KAPA Stranded mRNA-Seq Kit (Roche) according to the manufacturer’s standard protocol for mRNA capture, fragmentation, random-primed first strand synthesis, second strand synthesis with dUTP marking, A-tailing, adaptor ligation, and library amplification. Libraries were pooled and multiplex sequenced on an Illumina NovaSeqX Plus System (150bp, paired-end, >25 × 10^6^ reads/sample).

Adapter sequences were trimmed from raw reads using Cutadapt (v5.0) (error rate of 0.1, 3bp minimum overlap, minimum 3′ base quality score of 25, and a minimum post-trimming read length of 55 bp. Poly(A) tails and poly(G) sequences longer than 20 bp were also removed.

Trimmed reads were aligned to the Homo sapiens reference genome (GRCh38), incorporating exon, splice-site, and SNP annotations, using HISAT2 (v2.2.0). Resulting SAM files were converted to sorted BAM files using SAMtools (v1.21), and gene-level read counts were quantified with featureCounts (v2.0.6).

## Statistics

Gene counts were log2-transformed, and the 3,000 most variable genes, ranked by standard deviation, were selected for principal component analysis (PCA). Differential gene expression analysis between treatment groups was conducted using the edgeR package (v4.4.2) in R (v4.4.2). Low-expressed genes were filtered using the filterByExpr function with treatment- group as the experimental factor. Library sizes were recalculated after filtering and normalization performed using the trimmed mean of M-values (TMM) method to correct for sample-specific variations. Gene-wise dispersions were estimated, and a negative binomial generalized log-linear model was fitted to the normalized counts, incorporating an interaction term between sample type and treatment. Differential expression between treatment groups within each sample type was assessed using gene-wise quasi-F tests for the relevant model contrasts. Resulting p-values were adjusted for multiple testing using the Benjamini–Hochberg (BH) method, with adjusted p-value threshold of 0.05 for statistical significance. Gene set enrichment analysis and over- representation analysis were performed using clusterProfiler package (v4.14.6), and overlap between human disease signatures and assembloid differentially expressed genes was evaluated using a hypergeometric test.

## Results

### Study overview

Overview of our study is in Fig 1. Using the previously established, a transcriptomics-based computational drug repositioning pipeline,[15] we queried gene expression signatures of endometriosis against the CMAP database and identified 299 therapeutic candidates, including fenoprofen, simvastatin and primaquine. In our rat endometriosis model, fenoprofen, primaquine, and simvastatin were found to reduce vaginal hyperalgesia.[15] We used the same drugs on the individual endometrial cell lines and both types of assembloids. We then performed bulk RNA sequencing of the individual cell lines and assembloids to assess gene expression from treated and untreated tissues.

**Figure 1.**
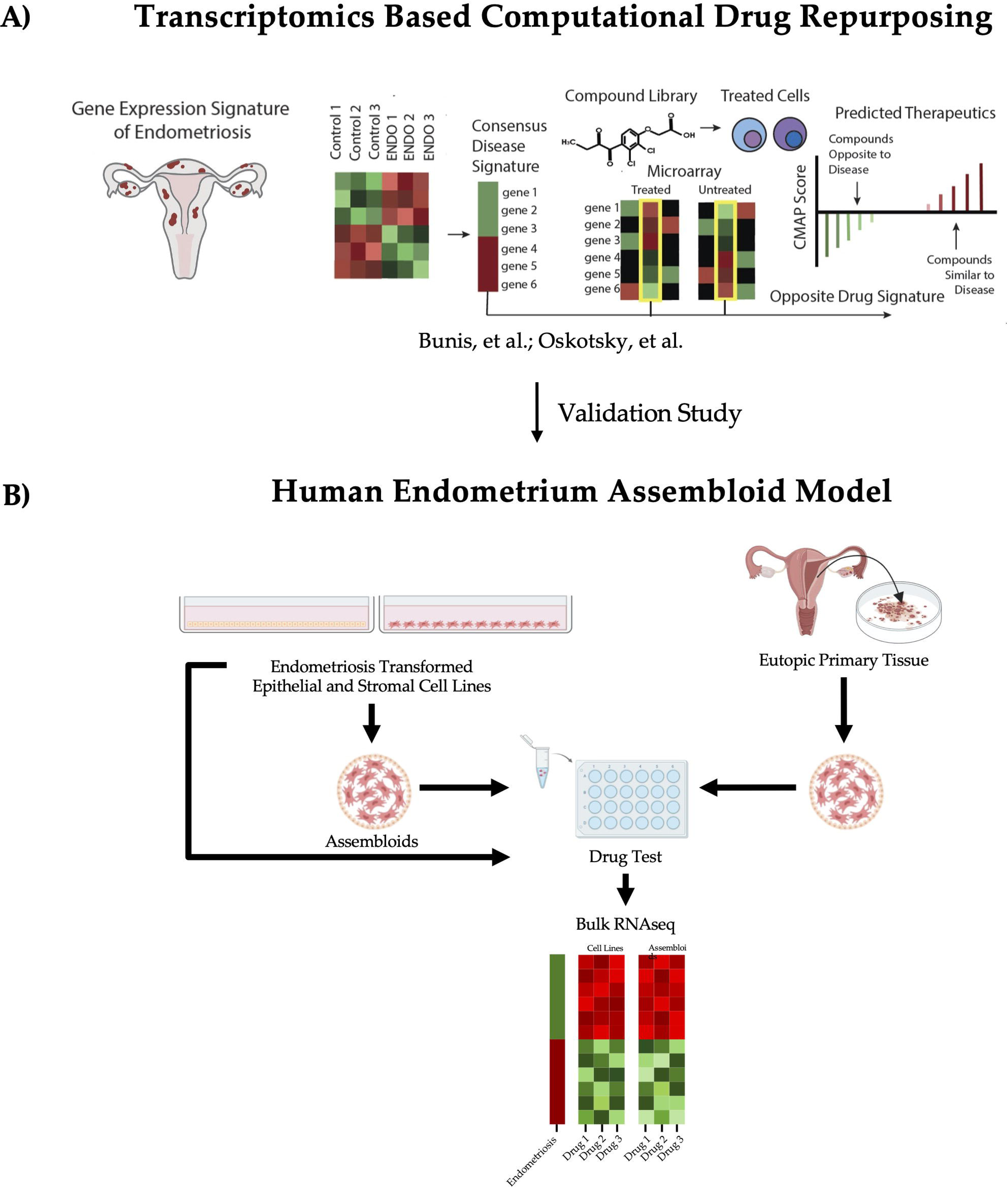
Study overview. (A) Transcriptomics Based Computational Drug Repurposing: An overview of the drug repositioning pipeline. First, gene expression signatures of endometriosis are queried against the Connectivity Map (CMAP) database containing over 1.5M gene expression profiles. We hypothesize that effective therapeutics can reverse disease-associated gene expression. 299 therapeutic candidates, including fenoprofen, simvastatin, and primaquine were identified. (B) Validation of therapeutic candidates using human endometrium assembloid model. Endometriosis transformed cell lines and eutopic tissue from endometriosis patients were used to produce the assembloids. We tested simvastatin, primaquine, ibuprofen, and fenoprofen on both the assembloids and individual epithelial and stomal cell lines. Bulk RNA sequencing was performed and the data was analyzed. Created in BioRender. Liu, J. (2026) https://BioRender.com/9vd2lop

### Establishment of assembloids from epithelial and stromal cell lines and eutopic endometrial tissue

Cell-line derived assembloids were created using a protocol adapted from Murphy et al (2019)[18]. Optimal epithelial-to-stromal cell ratios are not well-defined, largely due to challenges in accurate quantification caused by cell clumping in primary cultures. Established cell lines do not exhibit this limitation. Endometrial epithelial and stromal cells were therefore cultured separately and combined at varying ratios to determine the condition which yielded optimal assembloid formation. Increased proportions of stromal cells resulted in enhanced assembloid proliferation (Fig 2A, 2B). Based on these findings, the stromal cell proportion was increased to a 1:20 epithelial-to-stromal ratio for subsequent experiments, and the assembloids were analyzed by H&E staining. Resulting assembloids exhibited two distinct cellular layers with epithelial cells forming the outer layer and stromal cells forming the inner layer (Fig 2C). Using the same protocol, eutopic endometrial assembloids were established using cells obtained from patients with endometriosis. In contrast to cell line–derived assembloids, higher stromal cell counts were not correlated with better proliferation in patient-derived assembloids (Fig 2D). Once established, individual cell lines and the two types of assembloids were treated with fenoprofen, simvastatin, primaquine, ibuprofen (positive control), and vehicle (negative control) for 24 hr.

**Figure 2.**
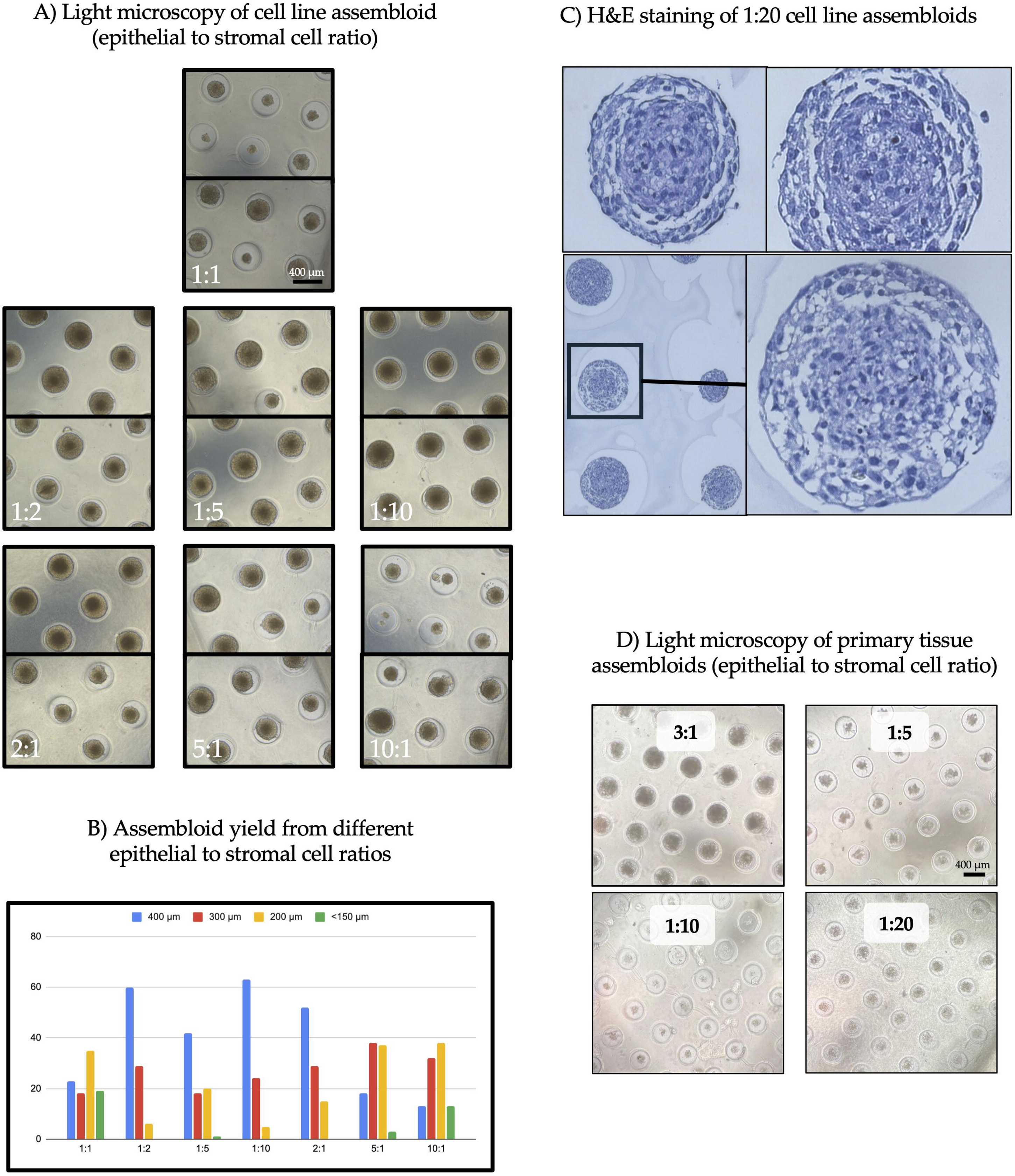
Assembloid development. (A) Light microscopy of endometriosis cell line assembloids. Cell line assembloids were created using endometrial epithelial and stromal cells at differing ratios. (B) Assembloid yields from different epithelial to stromal cell rations. More stromal cells produced larger assembloids. (C) H&E staining of 1:20 cell line assembloids. Lower left image is a magnified image of the lower right assembloid. Two distinct layers of cells can be seen. (D) Light microscopy of primary eutopic tissue assembloids. Higher epithelial cells to stromal cell ratio produced larger assembloids.

### RNA-seq analysis of drug treated cell lines and assembloids

To gauge the impact of candidate drugs on gene expression, bulk RNA sequencing was performed on the cell lines (N=3 for each cell type and drug condition), cell line assembloids (N=1 for each drug condition) and primary assembloids (N=2 for each drug condition).

#### Distinct transcriptional mechanisms underlying drug effects

Sample type had stronger effects on the transcriptomic profile than treatment, as distinct sample types formed separate clusters in the PCA plots (Supplementary Figure 1A-B). Specifically, cell line-derived assembloids, primary endometrial tissue assembloids, human endometrial epithelial cells, and stromal cells clustered separately, whereas treatments did not result in marked separation (Supplementary Figure 1A-B). For both epithelial and stromal cells, simvastatin-treated samples formed distinct clusters relative to other drug treatments, indicating simvastatin exerted strong and unique transcriptional effects in these endometrial cell types (Figure 1A-B, Supplementary Figure 1C-D).

We determined the differentially expressed genes (DEGs) induced by each treatment within each sample source (Figure 3C, Supplementary Figure 2). Simvastatin induced the most dramatic transcriptional perturbations with the highest numbers of DEGs (FDR < 0.05) across all sample types (nDEGs = 1147, 6531, and 5629 in assembloids, epithelial, and stromal cells, respectively, FDR< 0.05, Figure 3C). Primaquine showed a strong effect in epithelial cells with 4015 DEGs (FDR< 0.05) but demonstrated limited effects in stromal cells and primary endometrial tissue assembloids (365 DEGs in assembloids and 402 DEGs in stromal cells, FDR < 0.05). Surprisingly, the two NSAIDs, ibuprofen and fenoprofen, had minimal impact on cell lines and assembloids, with the fewest DEGs across all sample types compared to the other two drugs. This suggests that the analgesic effects of these NSAIDs are unlikely to be mediated through direct action on epithelial or stromal cells.

**Figure 3.**
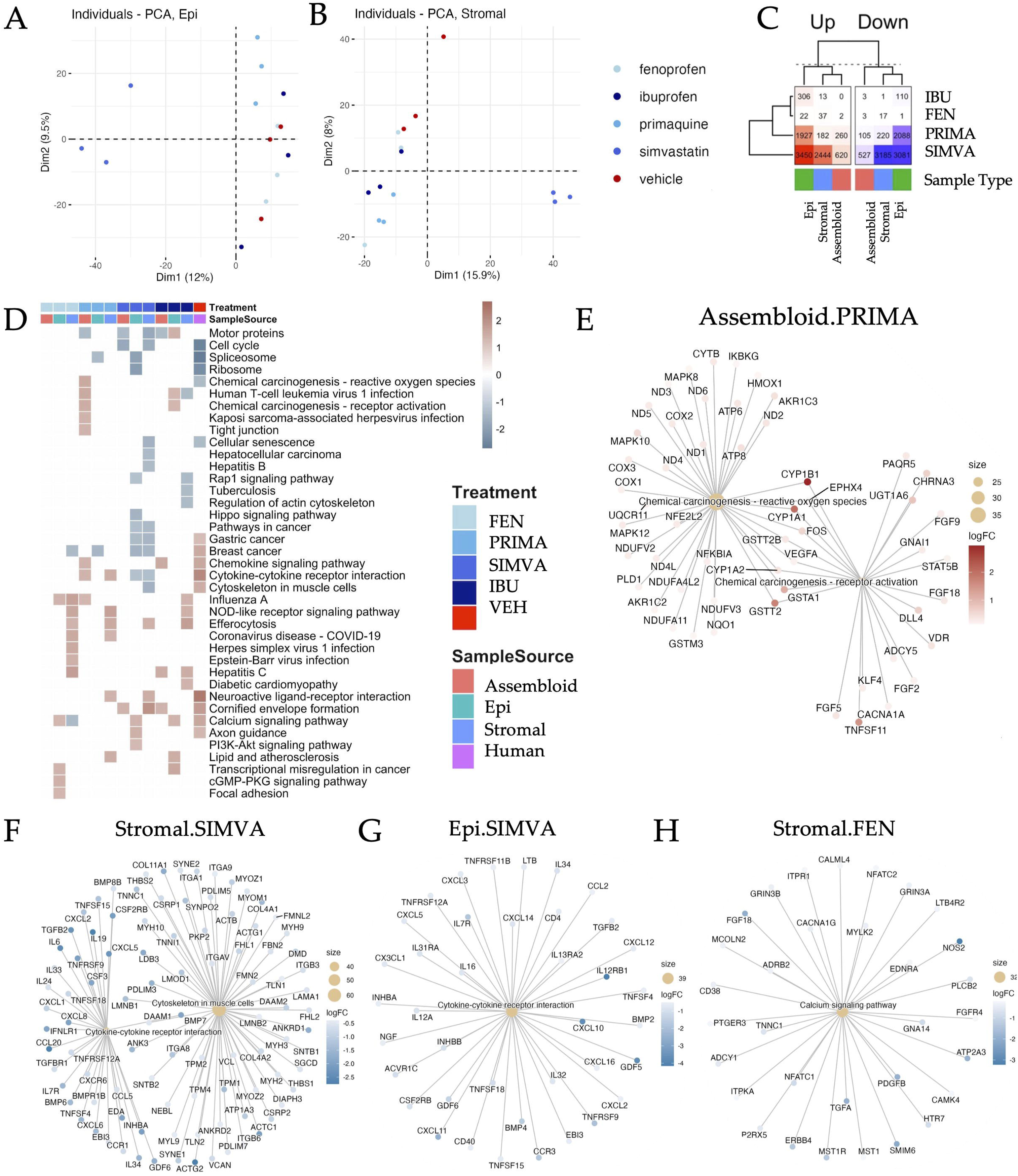
Transcriptional changes in cell lines and assembloids in response to treatments. (A) PCA plot of epithelial cells colored by treatment. (B) PCA plot of stromal cells colored by treatment. (C) Numbers of up- and down-regulated DEGs in assembloids, epithelial cells, and stromal cells exposed to candidate drugs. (D) Gene set enrichment analysis (GSEA) of KEGG pathway alterations in assembloids, epithelial cells, and stromal cells treated with candidate drugs, compared with endometriosis-associated transcriptional signatures derived from human endometrium samples. (E–H) Enriched pathways and core genes in endometrial assembloids treated with primaquine (E), in stromal cells treated with simvastatin (F), in epithelial cells treated with simvastatin (G), and in stromal cells treated with fenoprofen (H). In panels D-H, red indicates upregulated genes or pathways, and blue indicates downregulated genes or pathways.

To elucidate the mechanisms of these drugs across different sample types, we performed gene set enrichment analysis to identify biological pathways involved in drug-induced transcriptional changes and compared them to those altered in endometriosis, based on previously published microarray data from human eutopic endometrium (Figure 3D). The results revealed that primaquine reversed the chemical carcinogenesis-reactive oxygen species (ROS) pathway in endometriosis assembloids. Simvastatin reversed cytokine-cytokine receptor interaction in epithelial and stromal cell lines, and the cytoskeletal in muscle cells pathway in stromal cell lines. Fenoprofen reversed the calcium signaling pathway in stromal cell lines.

CYP1A1 and CYP1B1 were the two most highly upregulated genes within the chemical carcinogenesis pathway, upregulated in assembloids treated with primaquine, and are also key enzymes involved in estrogen biosynthesis (Figure 3E). In human endometrial data, CYP1A1 was over-expressed in endometrium from patients while CYP1B1 was downregulated.[15]

Simvastatin attenuated cytokine-cytokine receptor interaction by downregulating the expression of multiple interleukins and their receptors: IL16, IL12A, IL32, IL34, IL31RA, IL13RA2, and IL12RB1 (Figure 3F). Simvastatin reduced expression of several cytokine ligands: CXCL11, CX3CL1, CXCL5, CXCL3, CXCL14, CCL2, CXCL12, CXCL10, CXCL16, and CXCL2. Together, these findings suggest that therapeutic effects of simvastatin in endometrial cells may be mediated through suppression of pro-inflammatory responses, a central pathogenic feature of endometriosis.

In stromal cells, simvastatin also downregulated genes involved in cytoskeletal remodeling, a process that is critical for cell invasion (Figure 3G). Estrogen enhances cytoskeletal and membrane remodeling in endometrial stromal cells, resulting in enhanced cell motility and invasion in a variety of endometrial disorders.[20] In the human endometrial stromal cell lines, simvastatin treatment led to reduced expression of genes associated with actin nucleation and formin-mediated filament assembly, including DIAPH3, DAAM1, DAAM2, FMN2, and FMNL2, and genes encoding core cytoskeletal components such as ACTB, ACTG1, ACTC1, ACTG, MYH9, MYH10, MYH2, MYH3, MYL9, MYOM1, TPM1, TPM2, and TPM4 (Figure 3G). Also, genes involved in focal adhesion formation were also downregulated: ITGA1, ITGA8, ITGA9, ITGAV, ITGB3, ITGB6, TLN1, TLN2, and VCL (Figure 3G). These results suggest that enhanced cytoskeletal and membrane remodeling observed in endometriosis may be abrogated by simvastatin via its actions on stromal cells, potentially limiting their invasive ness.

The two NSAIDs reversed a few pathways altered in human eutopic endometrium. Ibuprofen, most commonly used for endometriosis pain, did not reverse any dysregulated pathways (Figure 3D). In contrast, in stromal cells fenoprofen suppressed calcium signaling and PTGER3, a pro-inflammatory gene upregulated in endometriosis, [21] and NFATC1, an inducer of COX-2 expression contributing to endometrial tissue proliferation[22] (Figure 3H).

## Discussion

This study established human endometrial assembloids using two endometriosis-based cell lines and primary cell lines from eutopic endometrial tissue of patients with endometriosis to test pathway reversals by simvastatin, primaquine, and fenoprofen identified in our drug-repositioning studies.[15] We found that simvastatin-treated samples formed distinct clusters from those treated with other drugs in both endometriosis-derived cell types on PCA plots. It also induced the highest number of DEGs across all sample types. Primaquine showed a strong effect in epithelial cells, suggesting its therapeutic effects may predominantly be mediated through epithelial-specific mechanisms. While fenoprofen did not produce substantial changes in the assembloids, it suppressed calcium signaling in stromal cells.

### Comparing assembloids to animal models

Our work serves as a valuable proof-of-concept for using human endometrial assembloids as a platform for drug testing potential endometriosis therapeutics.[23] The co-culture assembloid model of both epithelial and stromal cells in a 3D structure extends earlier epithelial-only organoid models.[24,25] This spheroid architecture, with epithelial cells surrounding encapsulated stromal cells, provides physiologically relevant tissue structure and microenvironment[26] and reproduces paracrine signaling that occurs between these two cell types in vivo. This human-centric approach provides an essential alternative to the systemic complexity and inherent biological limitations of animal studies.

### Rodent models have traditionally been used to study endometriosis

Immunocompromised mice with human endometrial tissue xenografts are often used to study disease establishment and progression.[27,28] In rats ectopic lesions undergo neoangiogenesis, leading to vaginal hyperalgesia.[29] Researchers, including our team, have used methods to measure changes in pain sensitivity and test efficacy of therapies.[16] Drug testing using rodent models of endometriosis are also increasing.[16,30] While animal models have a complete repertoire of cell types (epithelial, mesenchymal, endothelial, immune), and thus offer an orthogonal complement to findings with assembloids and organoids, rodents do not naturally menstruate and thus do not spontaneously develop the disease as humans do. This fundamental physiological divergence underscores the limitations of animal-based drug screening and highlights the importance of 3D assembloids, which preserve the human-specific hormonal responses and stromal-epithelial interactions necessary for authentic disease modeling.

#### Bulk RNA sequencing analysis of drug tested cell lines and assembloids

Bulk RNA sequencing analysis demonstrates the advantages of assessing drug effects in complex in vitro systems. Importantly, the drug response highly depended on sample source (cell lines vs. primary endometrial cell line assembloids vs. cell line assembloids), suggesting in vivo context is critical. Sample type had a stronger effect on the transcriptomic profile than the drug treatments, leading to distinct clusters on PCA plots. This highlights the risk of relying solely on single-cell monolayer cultures for therapeutic discovery, which may miss critical context-dependent responses observed in the 3D assembloid model. Our findings provide mechanistic evidence supporting use of simvastatin and primaquine as potential therapies for endometriosis.

#### Simvastatin and primaquine: potential treatments for endometriosis

Simvastatin has been previously considered as a potential treatment for endometriosis, with prior studies demonstrating its ability to inhibit human endometrial stromal cell proliferation and invasion.[31] In a rat model of endometriosis, simvastatin downregulated cytokine-cytokine receptor interaction signaling in lesions and eutopic uteri, and up-regulated cytoskeleton in muscle cell-related pathways in lesions.[16] In baboons, simvastatin also reverted serum levels of miR-150-5p and miR-451a, and miR-3613-5p toward control expression levels.[32] In the current study, simvastatin demonstrated the most significant transcriptional perturbations across all sample types. Mechanistically, simvastatin’s therapeutic effects occur via two primary actions on endometriosis-derived cells: suppression of proinflammatory features and inhibiting invasion. It reversed cytokine-cytokine receptor interaction in both epithelial and stromal cell lines. This involved downregulating key interleukins and their receptors (e.g., IL16, IL12A, IL32) and cytokine ligands (e.g., CXCL11, CXCL5, CCL2). Proinflammatory responses are central to endometriosis pathogenesis, thus suggesting a major therapeutic avenue.

Simvastatin also downregulated genes involved in cytoskeletal remodeling and focal adhesion formation in stromal cells (e.g., DIAPH3, ACTB, ITGA1, VCL), a process critical for cell motility and invasion. Simvastatin may thus limit endometriotic stromal cells capacity to invade surrounding tissues. Moreover, the anti-inflammatory and inhibition of cell motility/invasion properties of simvastatin are consistent with a role for it in inhibiting inflammatory pain and disease progression, respectively.

Primaquine showed a strong, epithelial-specific effect, reversing the chemical carcinogenesis-reactive oxygen species (ROS) pathway in endometriosis assembloids. It specifically upregulated the estrogen biosynthesis enzymes CYP1A1 and CYP1B1. CYP1A1 and CYP1B1 are isoforms of cytochrome P450 that catalyze the biotransformation of estrogens.[33] This suggests primaquine’s benefits may be predominantly mediated through epithelial-specific mechanisms, offering a more targeted approach.

#### Limitations

A key limitation of the current study is the small sample size, which was unavoidable due to the restricted number of patients who met the inclusion criteria for the study, specifically, those not on hormonal contraceptives. A key factor for interpreting the bulk RNA sequencing results is that the drug response was highly dependent on sample source (cell lines vs. primary tissue assembloids), as noted above.

## Conclusion

The endometrial assembloid model serves as a robust and relevant platform for drug testing. Simvastatin emerged as the most promising candidate, demonstrating both anti-inflammatory and anti-invasion mechanisms of action. These proof-of-concept experiments underscore the promise of assembloid models to test therapeutic candidates for endometriosis, providing a bridge between in silico discovery and in vivo validation. The application of this model can accelerate translation of repositioned and novel compounds toward clinical trials for this common and debilitating disease.

Supplementary Figure 1. Principal component analysis of samples colored by sample type and treatment. (A) PCA plot of all samples colored by sample types. (B) PCA plot of all samples colored by treatment. (C) PCA plot of cell line-derived assembloids colored by treatment. (D) PCA plot of primary endometrial tissue-derived assembloids colored by treatment.

Supplementary Figure 2. Volcano plots of differential gene expression profiles across sample types in response to drug treatments. The x-axis reflects (-) log10 of the Benjamini-Hochberg (BH)-adjusted p-value, and the y-axis reflects the log2 fold change.

Article Type: Laboratory based study

## Data availability

The RNA seq data are available through the Gene Expression Omnibus (GEO), accession ID GSE296883 (https://www.ncbi.nlm.nih.gov/geo/query/acc.cgi?acc=GSE296883).

CRediT Authorship Contribution Statement: FA (Conceptualization, data curation, formal analysis, investigation, methodology, writing – original draft), XT (Data curation, formal analysis, writing – original draft), JCI (Conceptualization, investigation, data curation, writing – review and editing), BL (writing – original draft), TTO (Conceptualization, writing – review and editing), MS (Conceptualization, funding acquisition, writing – review and editing, final approval of manuscript), FJM (Conceptualization, funding acquisition), LCG (Conceptualization, funding acquisition, writing – review and editing, supervision, and final approval of manuscript).

## Attestation statements

Data produced in the study has not been previously published. Data will be made available to the editors of the journal for review or query upon request.

## Capsule

This study tested drug-repositioning candidates using human endometrial assembloids. Simvastatin and primaquine reversed key disease-associated pathways, validating assembloids as a promising model for identifying effective, well-tolerated endometriosis treatments.

## Supporting information

Supplementary Figure 1. Principal component analysis of samples colored by sample type and treatment

Supplementary Figure 2. Differential gene expression profiles across sample types in response to drug treatments.

## Acknowledgments

UC Davis Comprehensive Cancer Center Genomics Shared Resource (NCI P30 CA93373)

