## Supplementary figures and images for "Simvastatin and Primaquine Identified as Potential Endometriosis Therapeutics via a Novel Epithelial-Stromal Assembloid Drug Screening Assay"

### Supplementary Figure 1. Principal component analysis of samples colored by sample type and treatment

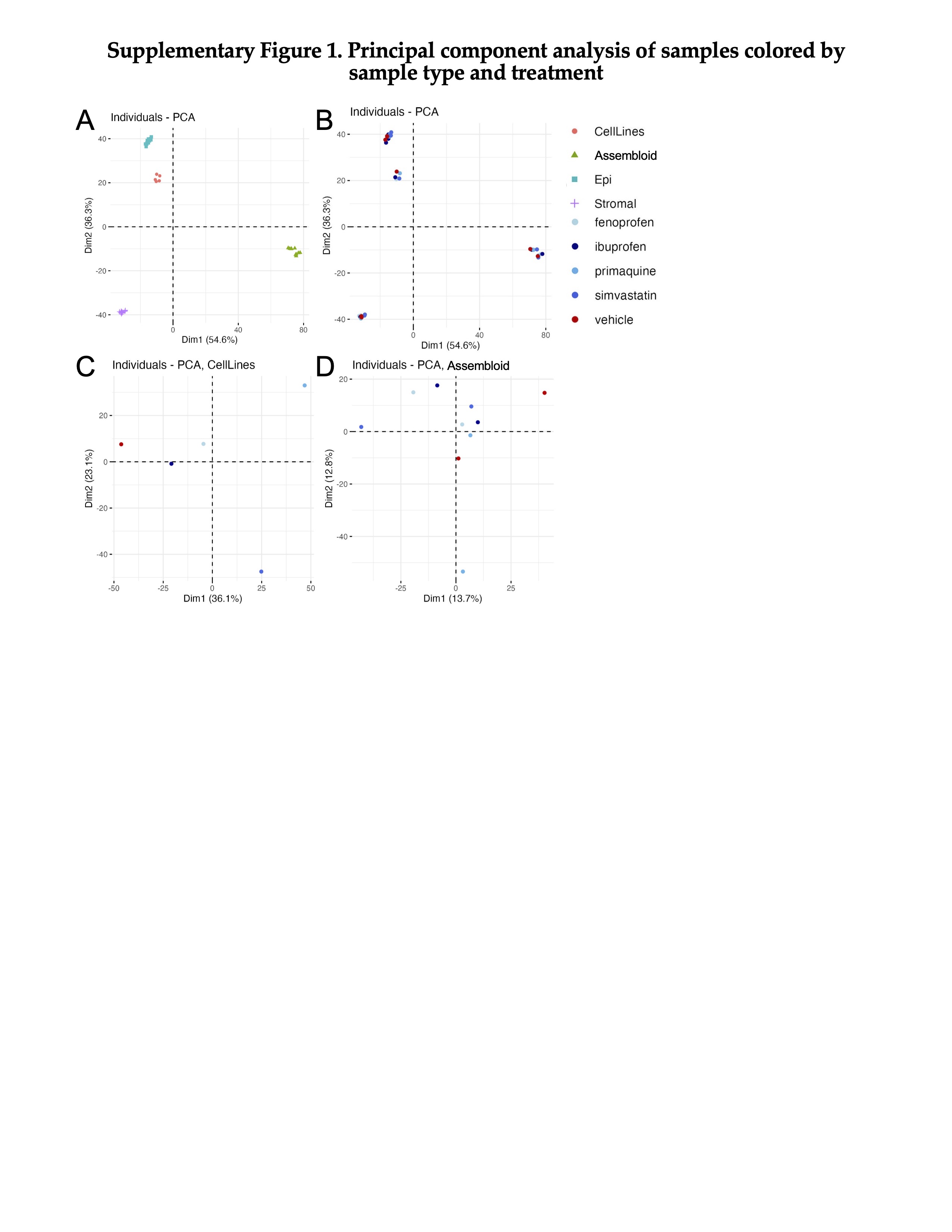

### Supplementary Figure 2. Differential gene expression profiles across sample types in response to drug treatments.

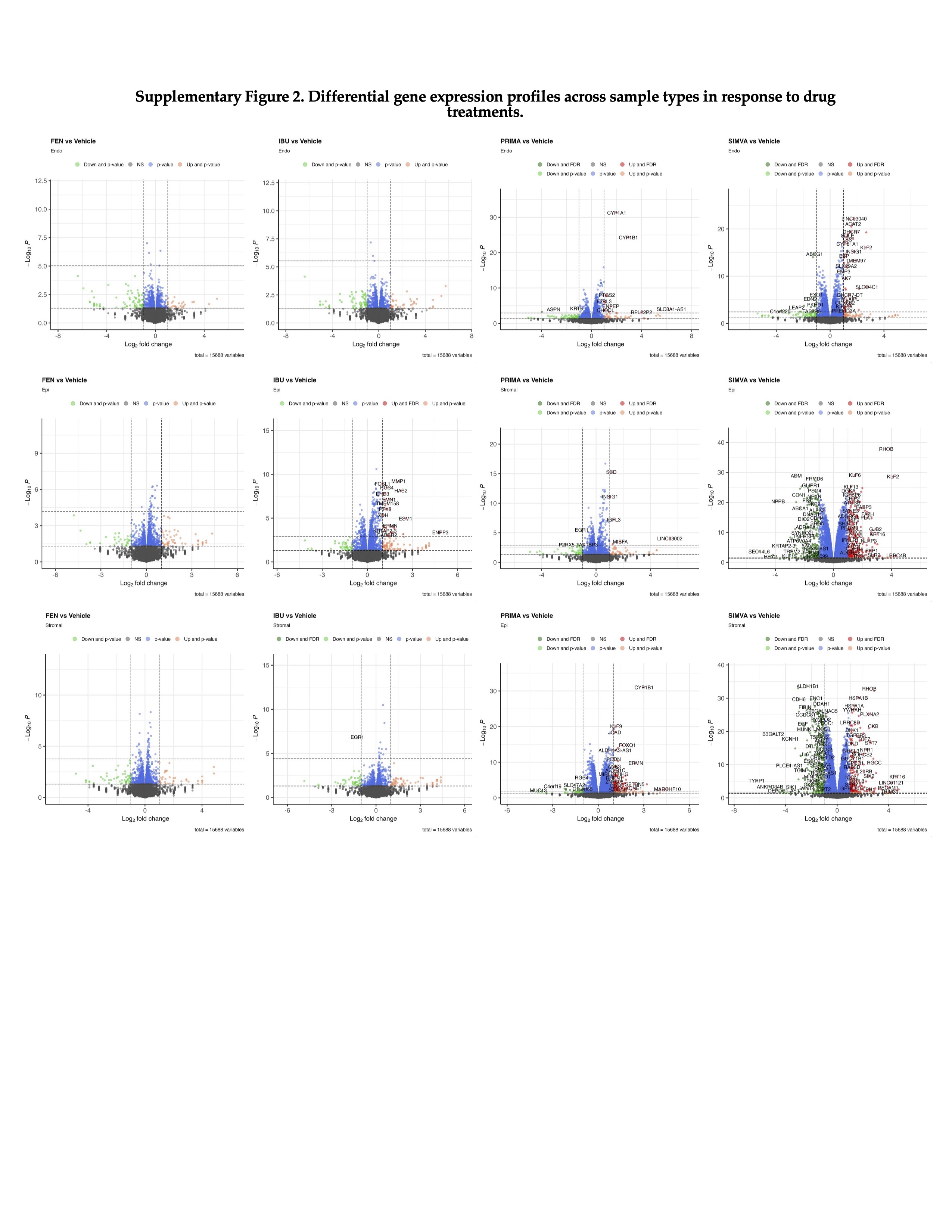
